# A role for the cortex in sleep-wake regulation

**DOI:** 10.1101/2020.03.17.996090

**Authors:** Lukas B. Krone, Tomoko Yamagata, Cristina Blanco-Duque, Mathilde C. C. Guillaumin, Martin C. Kahn, Colin J. Akerman, Anna Hoerder-Suabedissen, Zoltán Molnár, Vladyslav V. Vyazovskiy

## Abstract

The cortex and subcortical circuitry are thought to play distinct roles in the generation of sleep oscillations and global control of vigilance states. Here we silenced a subset of cortical layer 5 pyramidal and dentate gyrus granule cells in mice using a cell-specific ablation of the key t-SNARE protein SNAP25. We found a marked increase in wakefulness accompanied by a reduced rebound of EEG slow-wave activity after sleep deprivation. Our data illustrates an important role for the cortex in both global state control and sleep homeostasis.

## Main text

The duration, timing and architecture of sleep are strictly regulated. Early studies based on neurological case reports, transections and electrical stimulation suggested that global state transitions are mediated via a distributed circuitry across the brainstem, the hypothalamus and the basal forebrain^1,2^. More recent studies using selective targeting of specific neuronal populations based on their gene expression or connectivity patterns, highlighted that the sleep-wake promoting circuitry is highly complex, with distinct subcortical brain regions and neuronal subtypes responsible for specific aspects of wakefulness and sleep^3,4^. Although sleep-wake states are defined by the occurrence of neocortical and hippocampal oscillations^5^, the possibility that cortical structures control vigilance states has been overlooked.

Cortical oscillations and neuronal firing patterns mirror sleep homeostasis^6,7^. Sleep homeostasis refers to the adjustment of duration and intensity of sleep, to the duration of preceding wakefulness^6^. Electroencephalogram (EEG) slow-wave activity (SWA, EEG spectral power between 0.5-4 Hz) represents a reliable marker of sleep-wake history^6^ and has been proposed to underlie many functions of sleep, such as cellular maintenance^8^ and synaptic plasticity^9^. SWA can be regulated in a local, use-dependent manner^10,11^, in line with the view that sleep emerges within cortical networks driven by the local accumulation of metabolic products, such as adenosine^12^. However, slow waves also occur under anaesthesia, in isolated cortical slabs or even *ex vivo*^13–16^. Therefore, the capacity to produce slow waves does not automatically imply a causative role for the cortex in sleep or sleep homeostasis, either on a local or global level.

Here we test whether cortical structures have a function in regulating global sleep-wake dynamics. We focused on pyramidal neurons within layer 5 of cortex, which possess a set of potentially relevant properties including high dendritic spine density and axonal arborisation^17^, and high sensitivity to adenosine^18^. Furthermore, layer 5 pyramidal neurons, and specifically slow intrinsically bursting neurons unique to this layer^19^, have been shown to play a key role in the initiation and propagation of cortical slow waves^19–22^.

Laminar local field potential (LFP) and multiunit activity (MUA) recordings were performed from the primary motor cortex, concomitantly with EEG and electromyography (EMG) monitoring in male adult mice during 24 h baseline, as well as during and after 6 h of sleep deprivation (Figure 1a-d). As expected, the laminar profile of LFPs and MUAs revealed generally activated patterns during waking and rapid eye movement (REM) sleep. Correspondingly, during non-rapid eye movement (NREM) sleep, we observed depth positive LFP slow waves associated with a generalised suppression of spiking activity across cortical layers (Figure 1c)^23,24^. Neurons were generally more active in the deep layers of the motor cortex, especially in layer 5 (Figure 1c,e), in line with previous results^22^. Consistent with the idea of an active role for layer 5 in shaping sleep oscillations^17,20,21^, we also found that neurons in layer 5 tended to initiate spiking upon the onset of population ON periods (Figure 1e-g). A leading role for layer 5 was indicated by a stronger initial surge of neuronal firing at OFF-ON transitions (Figure 1f), and a shorter latency to the first spike during ON periods, even when the total number of spikes in layer 5 was matched with layer 2/3 (Figure 1g).

**Figure 1:**
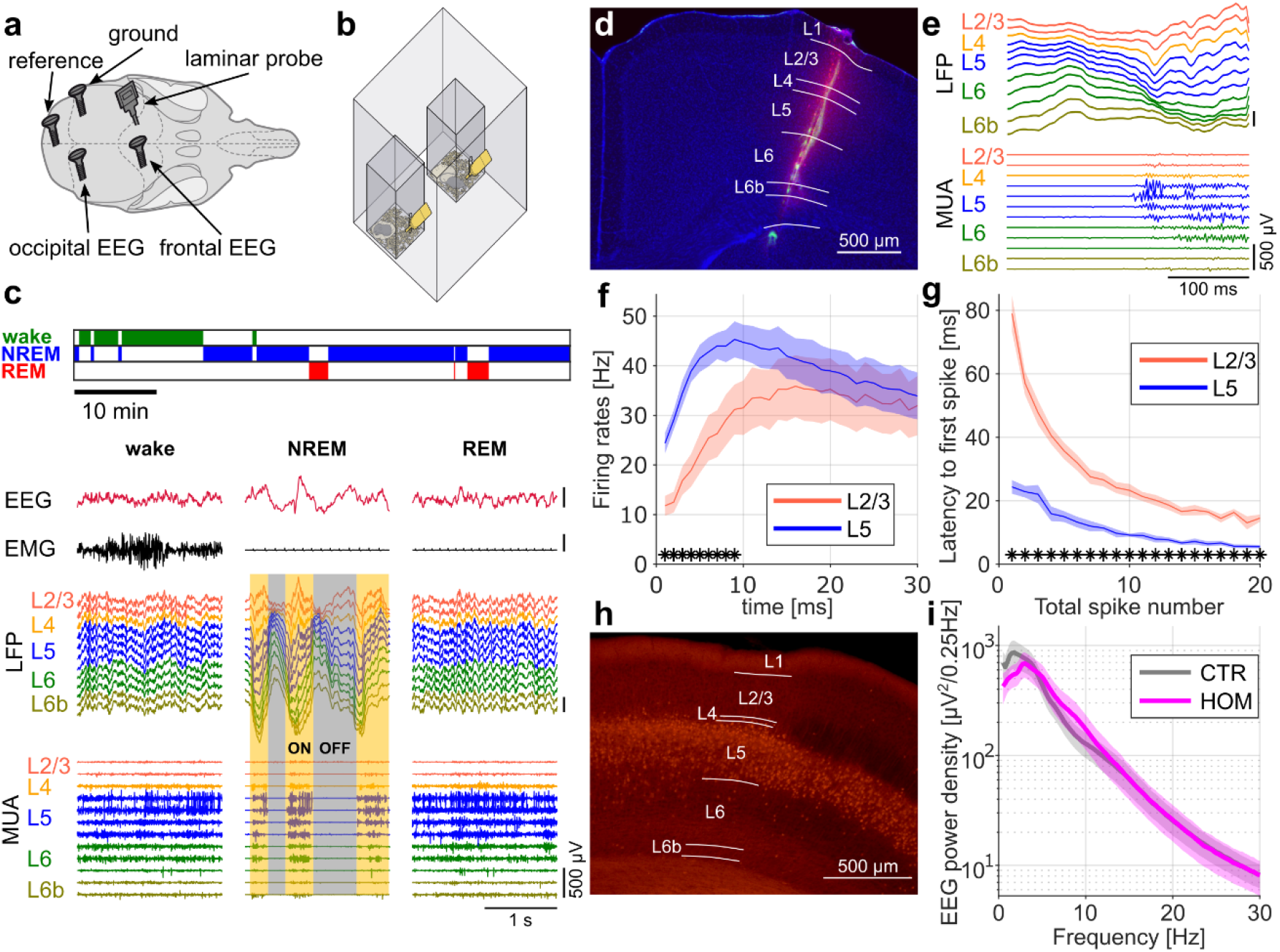
Cortical recordings in freely moving mice implicate layer 5 in sleep. a) Schematic of electrode positions in relation to the mouse skull. EMG electrodes in the neck muscles not depicted. b) Schematic of a Faraday chamber with two plexiglas cages for recording of individually housed mice. c) Representative hypnogram colour coded for the three vigilance states (wake, NREM, and REM sleep) together with EEG, EMG, LFP, and MUA traces showing characteristic signatures of the respective vigilance state. Neuronal ON and OFF periods during NREM sleep are highlighted with transparent yellow and grey panels, respectively. All scale bars on y-axis, 500μV. d) Representative histology of a laminar implant. The DiI trace (red) demarcates the electrode insertion tract while fine electrode marks (green) reveal the final depth of several individual channels on a DAPI counterstained (blue) coronal section. Microlesions of selected channels aid the classification of the recording channels into the cortical layers. Merged image using three fluorescent channels. e) Representative OFF-ON transition illustrating that population activity typically starts in layer 5. Both scale bars on y-axis, 500μV. f) Average firing rates at OFF-ON transitions. Note that layer 5 presents an earlier surge in firing rate compared to layer 2/3 (two-tailed paired t-test for each 1-ms time bin). g) Latency to the first spike for matched spike numbers during the first 200 ms of the ON period. Note that layer 5 has a shorter latency to the first spike compared to layer 2/3 for any given total spike count analysed (1 to 20 counts, two-tailed paired t-test for each total spike count). h) Representative histology of neocortical Cre-expression under the Rbp4-Cre promotor. Note that Cre-positive cells (red) in neocortex are largely restricted to layer 5. i) Frontal EEG spectra of homozygous cortical SNAP25-ablated (HOM) and control animals (CTR). Note that for none of the frequency bins a statistical difference was observed between HOM and CTR mice (two-tailed t-tests for independent samples performed on log-transformed spectra). N=7 for laminar analysis in WT animals (panels f,g). N=5 (CTR) and n=8 (HOM) for absolute EEG spectra of the cortical SNAP25-ablated mouse model (panel i). CTR: control animals. DiI: 1,1’-Dioctadecyl-3,3,3’,3’-Tetramethylindocarbocyanine Perchlorate. EEG: electroencephalogram. EMG: electromyogram. HOM: homozygous animals. LFP: local field potentials. MUA: multi unit activity. NREM: non-rapid eye movement sleep. REM: rapid eye movement sleep. WT: wild type animals.

To induce a cortex-wide reduction in the output from layer 5 pyramidal neurons, we generated a transgenic mouse line, in which a subpopulation (~ 15-30 %) of pyramidal cells in layer 5 of the neocortex lacked the key t-SNARE protein SNAP25 (Rbp4-Cre;Ai14;Snap25^fl/fl^)^25^. While Rbp4-Cre is known as a pan layer 5 driver line^26^, it additionally presents a strong Cre-expression in hippocampal dentate gyrus. By contrast, the Rbp4-Cre line is only very sparsely expressed in the hypothalamus (Suppl. Figure 1), a key brain area classically considered in the regulation of sleep-wake state transitions. Selective targeting of Rbp4-Cre to layer 5 was confirmed using DAPI histology to define cortical layers and a tdTomato reporter to identify Cre-expressing cells (Figure 1h and Suppl. Figure 2). Cre-dependent excision of exon 5a/5b leads to a reduced length gene transcript and non-detectable levels of SNAP25 protein in Cre-positive neurons^25^. Ablation of SNAP25 virtually abolishes calcium-evoked neurotransmitter release from the neurons carrying the mutation^25,27^, rendering the cells functionally silent. Transgenic mice (Rbp4-Cre;Ai14;Snap25^fl/fl^) did not show obvious physical, neurological or behavioural abnormalities, but their body weight was lower compared to Cre-negative controls (SNAP25-ablated: 20.9±0.6 g, controls: 23.4±0.6 g, t(13)=2.87, p=0.013, two-tailed t-test for independent samples), as reported previously^25^. The electrophysiological signals revealed typical signatures of wakefulness and sleep states in both genotypes, and, importantly, ablation of SNAP25 did not result in significant changes in absolute EEG power spectra during NREM sleep, or in the occurrence of slow waves (Figure 1i), but did show a leftward shift of the theta-peak frequency during REM sleep (Suppl. Figure 3). This result is in line with the observation that Rbp4 expression is not restricted to layer 5 of the neocortex, but is also present in granule cells of the hippocampal dentate gyrus^25^, which is involved in generating theta activity during REM sleep^28,29^.

While baseline differences in sleep oscillations between genotypes were modest, the overall daily sleep-wake profile of cortical SNAP25-ablated animals showed marked differences (Figure 2a). Control animals generally showed a higher amount of waking during the dark period as compared to the light period, but short bouts of sleep were common during the dark period, as typically observed in C57BL6 mice (Figure 2b)^30^. In contrast, cortical SNAP25-ablated animals showed unusually long wake bouts upon dark onset that often persisted for several hours or more (Figure 2b,e). On average, over 24h these animals spent 13.83±0.39 h awake, which is 31 % more than control animals (10.57±0.42 h, t(13)=5.55, p<0.001, two-tailed t-test for independent samples), and the amount of sleep decreased proportionally (Figure 2c). Interestingly, the differences between genotypes were primarily evident in the dark period (Figure 2d and Suppl. Figure 4), which is the mouse’s circadian active period and when the homeostatic sleep drive typically builds up to high levels as a result of prolonged wakefulness^30,31^. The increased capacity of cortical SNAP25-ablated animals to engage in prolonged wake episodes during the dark period suggests a potential attenuation of homeostatic sleep processes. To assess the build-up of the homeostatic sleep drive during wakefulness, we detected all consolidated wake bouts longer than 15 min across 24 h and compared the levels of EEG SWA during NREM sleep for 15-min time windows preceding and following individual wake episodes. As expected^31^, a positive correlation was observed in both genotypes, whereby longer spontaneous wake episodes were followed by proportionally higher levels of SWA during NREM sleep (Figure 2f). However, the increase of SWA relative to the duration of wake episodes was significantly smaller in cortical SNAP25-ablated animals compared to controls (Figure 2f). This finding indicates that ablation of SNAP25 in the neocortex and hippocampus affects the relationship between sleep-wake history and the levels of SWA.

**Figure 2:**
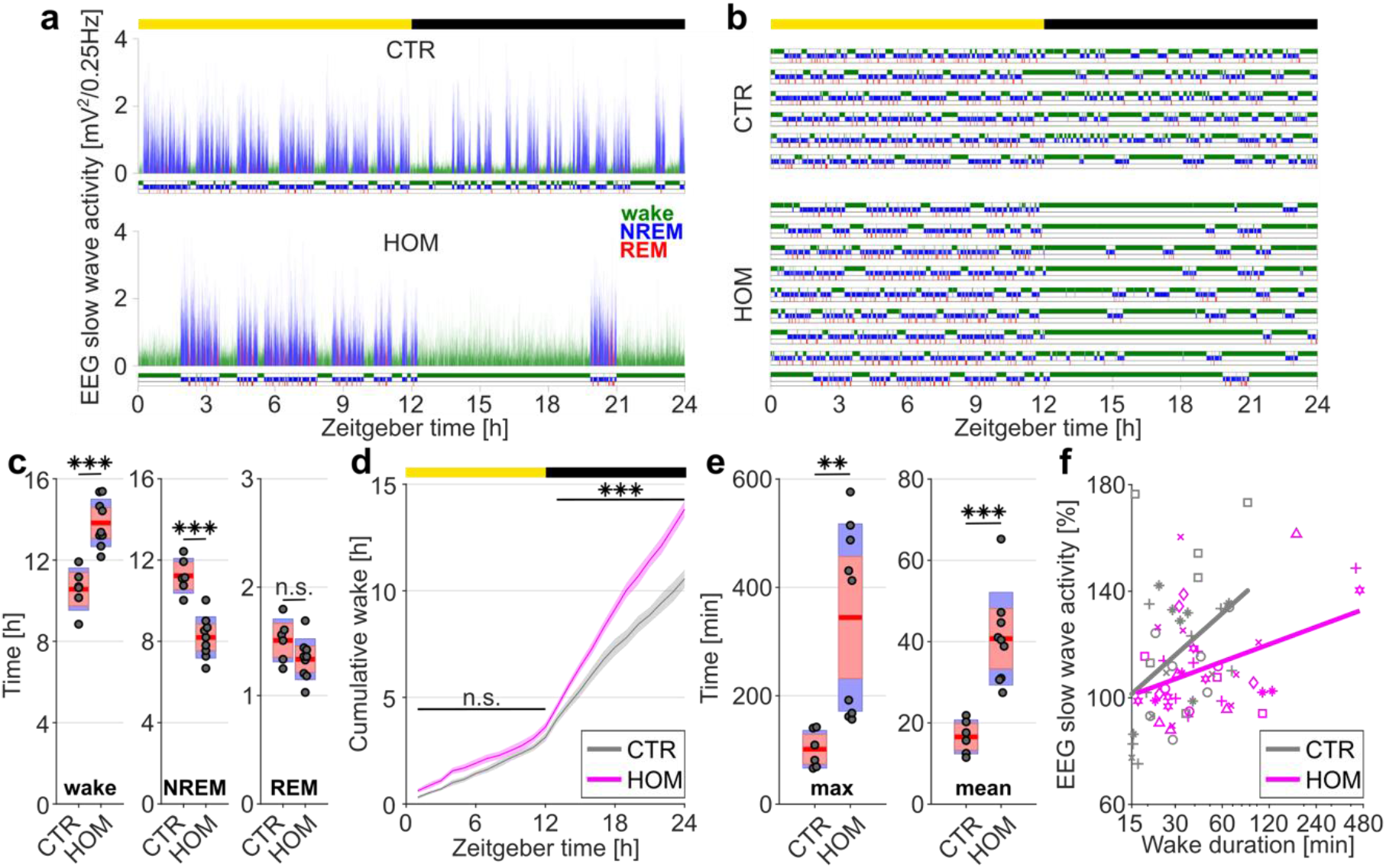
Selective cortical SNAP25-ablation alters sleep architecture. a) Hypnogram and EEG slow wave activity (0.5- 4.0Hz, 4s epochs) of one representative animal from each genotype in undisturbed 24h baseline recordings. b) Hypnograms from all individual control (CTR) and cortical SNAP25-ablated (HOM) animals under undisturbed baseline conditions. Note the increased amount of wakefulness and long wake episodes in HOMs. Vigilance states in A&B colour coded: wake=green, NREM=blue, REM=red. c) Time spent in vigilance states (wake, NREM, and REM) during 24h baseline recordings. Note that HOMs spent 3h and 16min more time awake compared to CTRs (t-tests for independent samples, wake: t(13)=−5.550, p<0.001, NREM: t(13)=6.052, p<0.001, REM: t(13)=1.724, p=0.108). d) Cumulative amount of wakefulness over 24h baseline recordings. Note that the slope of wake accumulation is similar between genotypes during the light period but steeper for HOMs compared to CTRs during the dark period (ZT12-24)(mixed ANOVA, within-subject factor ‘Light-Dark-Period’, between-subject factor ‘genotype’, main effect: F(1,13)=39.96, p<0.001. Post-hoc two-tailed t-tests for independent samples: light period: t(13)=−1.62, p=0.13; dark period: t(13)=−7.07, p<0.001). e) Maximum and mean duration of individual wake episodes over 42 hours (BL and SD day excluding the 6 h sleep deprivation period). Note that for HOMs both the maximum and mean duration of wake episodes is longer than for CTRs (t-test for independent samples. Maximum duration: t(8.94)=−4.11, p=0.003. Mean duration: t(13)=−4.94, p<0.001). f) Relationship between wake duration and relative SWA power during NREM epochs. Note the reduced slope of the relationship between SWA ratio and duration of wake episodes in HOMs (general linear model with 5 factors: mouseID, mouseID x episode duration (random factors), genotype, episode duration, and genotype x episode duration (fixed factors); dependent variable: SWA ratio post/pre sleep; F(1,57.37)=6.44, p=0.014). Individual animals are represented with different symbols. Number of animals: n=6 CTR and n=9 HOM for vigilance state analyses (panels c, d, e), n=5 CTR and n=8 HOM for analysis of SWA (panel f). Yellow and black bars above panels a, b, and d indicate light and dark periods, respectively. Panels c and e represent grouped data including group mean (red line), 95% confidence interval (pink box), and one standard deviation (blue box) with individual data points overlaid. BL: baseline. CTR: control animals. EEG: electroencephalogram. HOM: homozygous animals. NREM: non-rapid eye movement sleep. REM: rapid eye movement sleep. SD: sleep deprivation. ZT: zeitgeber time.

An established approach to investigate the dynamics of sleep homeostasis is sleep deprivation, which is typically performed starting at light onset, when mice in laboratory conditions usually sleep^7,30^. Sleep deprivation was successful in both genotypes, as only a minimal amount of time (SNAP25-ablated: 1.84±0.75 %, controls: 1.39±0.32 %, p>0.05, Mann-Whitney U test) was spent asleep during the 6-h interval when the mice were kept awake by providing novel objects. Typically, sleep deprivation leads to a subsequent increase in sleep intensity, indicated by increased SWA during NREM sleep and an increase in the amount of sleep, specifically NREM sleep^6,30^. However, we observed a striking difference in this homeostatic rebound between genotypes (Figure 3). Cortical SNAP25-ablated animals presented a marked attenuation of the initial increase of SWA during NREM sleep after sleep deprivation (relative SWA in SNAP25-ablated: 136.77±3.98 %, controls: 180.57±5.13 %, first 30 min after sleep deprivation, t(11)=6.78, p<0.001, two-tailed t-test for independent samples, Figure 3b). This attenuation was especially pronounced in frequencies between 1.5 and 3.75 Hz (Figure 3c). Interestingly, the difference was restricted to the frontal EEG but was not found in the occipital derivation, consistent with the idea that homeostatic elevation of SWA is a predominantly frontal phenomenon (Suppl. Figure 5)^32^. Since consolidation and fragmentation of sleep are sensitive behavioural measures of homeostatic sleep pressure^33^, we next analysed sleep architecture in greater detail. Both genotypes showed an increase in NREM sleep amount during the recovery period (Figure 3d), and the duration of NREM sleep episodes was initially longer than during baseline (Figure 3e). However, during the dark period following sleep deprivation, cortical SNAP25-ablated mice still presented markedly prolonged episodes of wakefulness and an overall reduced amount of sleep compared to controls, as if sleep pressure were low (Figure 3f,g). Taken together, the data support the conclusion that preceding sleep-wake history is not effectively encoded in cortical SNAP25-ablated mice.

**Figure 3:**
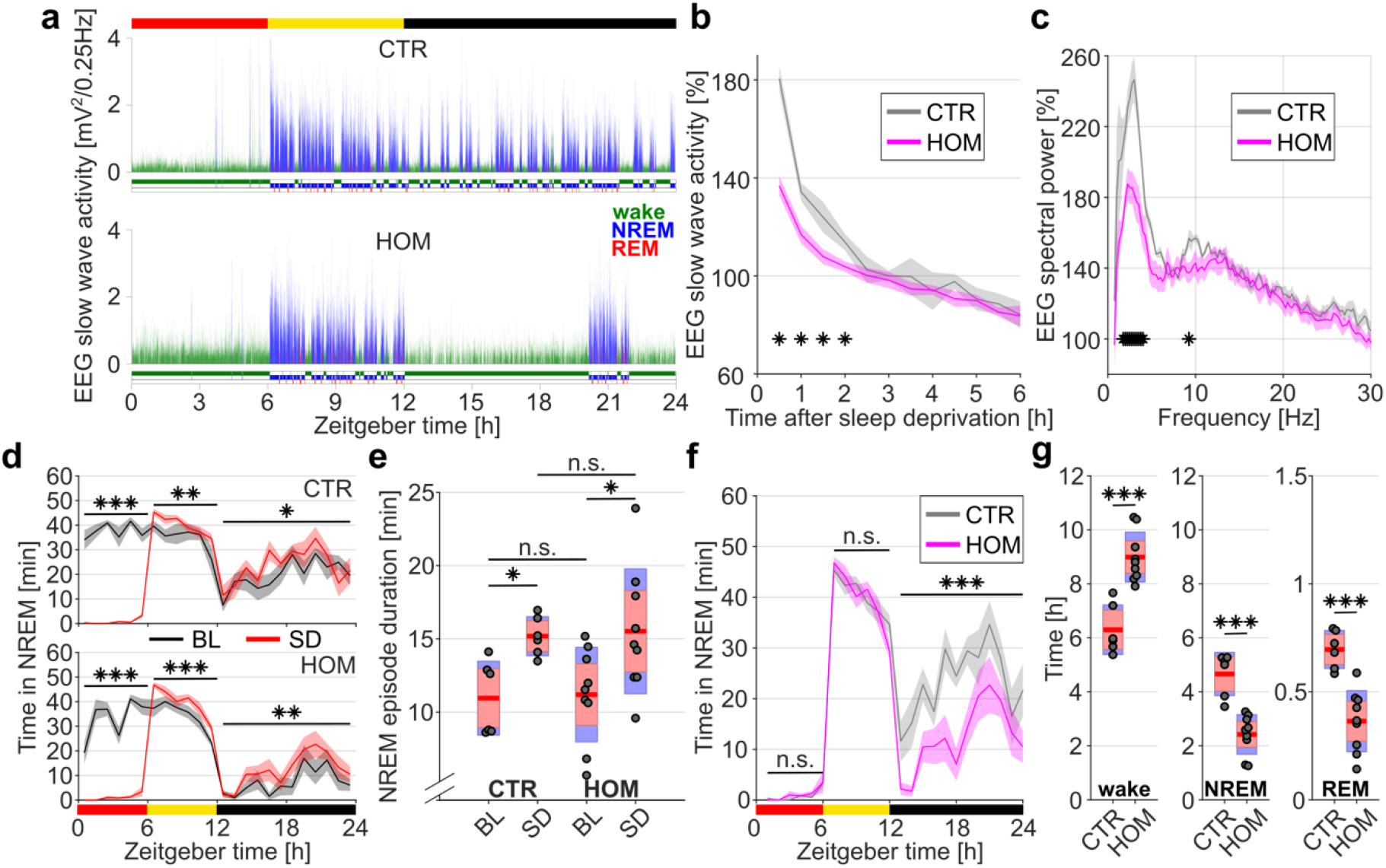
Selective cortical SNAP25-ablation alters homeostatic sleep regulation. a) Hypnogram and EEG slow wave activity (0.5-4.0Hz, 4s epochs) of one representative animal from each genotype in 24h recordings with 6h sleep deprivation (red bar above graph). b) Time course of EEG slow wave activity during NREM sleep after sleep deprivation relative to baseline average. Note that during the first 2 hours after sleep deprivation HOM animals show lower levels of SWA compared to CTR animals (mixed ANOVA, within-subject factor ‘time’, between-subject factor ‘genotype’, main effect: F(11,121)=7.65, p<0.001. Post-hoc comparisons with two-tailed t-tests for independent samples). c) EEG spectral power of NREM sleep during the first hour after sleep deprivation relative to baseline average. Note that the characteristic power increase in low frequencies following sleep deprivation is smaller in HOMs compared to CTRs (bin-wise comparison, frequency bins: 0.25 Hz, two-tailed t-tests for independent samples. Significant frequency bins: 1.5-3.75Hz and 9Hz). d) Time course of NREM sleep for each genotype compared between BL and SD days. Note the increase in NREM sleep compared to the respective baselines in both genotypes following sleep deprivation (red bar below graph). (repeated measures ANOVAs, within-subject factors ‘time’ and ‘condition’. Main effects: F(1.10,5.50)=184.84, p<0.001) for CTR and F(2,16)=195.24, p<0.001 for HOM. Post-hoc comparison with two-tailed t-tests for paired samples). e) Mean duration of NREM episodes during ZT6-7 on BL and SD days. Note that both genotypes show an increased duration of NREM episodes compared to their own baseline with no difference between genotypes (mixed ANOVA, within-subject factor ‘time’, between-subject factor ‘genotype’. No main effect for the interaction ‘time’ and ‘genotype’ or for between-subject factor ‘genotype’. Significant effect for within-subject factor ‘time’: F(1,13)=13.71, p=0.003. Post-hoc comparison of time windows in each genotype analysed using two-tailed t-tests for paired samples). f) Time course of NREM sleep on a sleep deprivation day compared between genotypes. Note that HOMs and CTRs do not differ in the time spent asleep during sleep deprivation (red bar below graph) or in the initial rebound sleep but HOMs sleep less during the 12 h dark period following sleep deprivation (mixed ANOVA, within-subject factor ‘time’, between-subject factor ‘genotype’. Main effect: F(1.16,15.01)=30.93, p<0.001. Post-hoc comparisons with two-tailed t-tests for independent samples). g) Time spent in vigilance states (wake, NREM, and REM) during the 12h dark period following sleep deprivation. Note that HOMs spent more time awake and less time asleep compared to CTRs (Mann-Whitney U test, p<0.001 for all vigilance states). N=6 CTR and n=9 HOM for vigilance state analyses (panels d, e, f, g), n=5 CTR and n=8 HOM for analysis of SWA and EEG spectra (panel b,c). Red, yellow and black bars above panels a, d, and f indicate sleep deprivation, light and dark periods, respectively. Panels e and g represent grouped data including group mean (red line), 95% confidence interval (pink box), and one standard deviation (blue box) with individual data points overlaid. The time course of NREM sleep in panels d and f is plotted in 1-h intervals. BL: baseline. CTR: control animals. EEG: electroencephalogram. HOM: homozygous animals. NREM: non-rapid eye movement sleep. REM: rapid eye movement sleep. SD: sleep deprivation. ZT: zeitgeber time.

To resolve the enigmatic functions of sleep, it is crucial to understand where and in what form the need to sleep is encoded in mammals, and how it is translated into an adequate compensatory response^34^. Our study reveals a novel role for the cortex in sleep-wake regulation. We show that cortical structures are not only responsible for the generation of state-specific oscillations, but also exert active control over sleep homeostasis and the global regulation of sleep-wake states. This finding supports the hypothesis that brain structures fundamentally involved in sleep regulation extend far beyond the traditionally considered subcortical circuitry^4,12,35^. The Rbp4-Cre mouse line is widely used as a driver line for neocortical layer 5 pyramidal neurons^20,26^, and altered neurotransmission in these cells is the most plausible cause for the striking sleep phenotype described here. In addition, we identify a potential contribution of archicortical dentate gyrus cells of the hippocampus. Given the critical place of the hippocampus in brain-wide circuitry involved in memory and temporal processing^36^, this structure may have a so far unrecognised role in encoding time spent awake or asleep. Therefore, whilst we cannot entirely exclude the role of developmental compensation or the possibility that a small proportion of cells were functionally silenced in other brain regions, the most parsimonious explanation is that cortical structures play a fundamental role in sleep regulation and actively contribute to the flipping of the sleep-wake switch.

Our results support the possibility that cortical structures generate sleep drive locally, in an activity-dependent fashion^8,12^. However, the data also support a novel role of the cortex in sensing and integrating signals of sleep need^12,37^. Intracellular processes reflecting wake-dependent increases in sleep need, conserved both in mammalian and non-mammalian species, may represent changes in synaptic phosphoproteome^38,39^, endoplasmic reticulum stress^40^, or redox homeostasis^41^. Extracellular signals may be found in molecular regulators of inflammation and plasticity^12^, or adenosine levels regulated through neural-glial interactions^37^. Arguably, such changes occur not only locally, but also across brain-wide distributed networks and must be combined to elicit a global homeostatic response, reflected in an occurrence of intense sleep, characterised by elevated cortical SWA and increased sleep propensity. We propose that the wide connectivity of layer 5 pyramidal neurons to other parts of cortex, thalamus, and potentially sleep-wake regulating nuclei in the brainstem^26^, places this neuronal population in an ideal position not only to generate SWA, but also to sense and integrate the signals related to sleep need, and ultimately broadcast the information to the subcortical circuitry responsible for sleep-wake switching^3^.

## Methods

### Animals

EEG/EMG implants were performed in 7 wild type C57BL/6 mice (WT, surgery weight 29.5±0.8 g, age at day of baseline recording 125±8 days), 9 Rbp4-Cre;Ai14;Snap25^fl/fl^ mice (HOM, surgery weight 20.9±0.6 g, age at day of baseline recording 86±6 days) and 6 Cre-negative littermates (CTR, surgery weight 23.4±0.6 g, age at day of baseline recording 78±5 days) under isoflurane anaesthesia as described previously^42^. For technical reasons, the EEG datasets from 1 HOM and 1 CTR had to be excluded from the analyses of EEG spectra, but were included in all other analyses. We recorded laminar LFP and MUA signals in WT mice across cortical layers in primary motor cortex (+ 1.1 mm AP (anterior), −1.75 mm ML (left), tilt −15° (left)) using 16-channel silicon probes (NeuroNexus Technologies Inc., Ann Arbor, MI, USA; model: A1×16-3mm-100-703-Z16) with a spacing of 100 μm between individual channels. In a subset of HOM and CTR animals, laminar recordings were also obtained but were not included in the manuscript due to the low number of animals.

Rbp4-Cre;Ai14;Snap25^fl/fl^ is a triple transgenic mouse line, which was designed as a model for functional silencing of cortical layer 5 pyramidal and dentate gyrus granule cells. Snap25^fl/fl^ is a transgene, with lox-P sites flanking the alternatively spliced exons 5a and 5b of the t-SNARE (target membrane soluble N-ethylmaleimide-sensitive factor attachment protein (SNAP) receptor) gene *Snap25* leading to virtually complete absence of SNAP25 and cessation of Ca^2+^-dependent evoked synaptic vesicle release^25^.

All mice were housed individually in open cages before surgery and in individually ventilated cages during a recovery period of about one week after surgery. For sleep recordings, mice were transferred to separate custom-made plexiglas cages (20.3 × 32 × 35 cm), which were placed in sound-attenuated and light-controlled Faraday chambers (Campden Instruments, Loughborough, UK), with each chamber fitting two cages. Animals were allowed free access to food pellets and water at all times and underwent daily health inspection. A 12 h:12 h light-dark cycle (lights on at 9 am, light levels 120–180 lux) was implemented and temperature maintained at around 22 ± 2°C.

### Experimental procedures

After an acclimatization period of at least 3 days during which animals were habituated to the tethered recording conditions, a 24 h period of continuous recording starting at light onset was performed on a designated baseline day. On the subsequent day, all animals were sleep deprived for 6 h starting at light onset. Sleep deprivation was performed during the circadian period when mice are typically asleep and thus the homeostatic response to sleep loss can be most reliably elicited^30^. At light onset, recording chambers were opened, the nesting material removed, and novel objects placed into the mouse cages to encourage exploratory behaviour. Experimenters continuously observed the mice and exchanged the provided objects for new objects when mice stopped exploring. At the end of the 6 h sleep deprivation, all objects were removed, the nesting material returned, and the recording chambers closed.

### Data acquisition, data processing, and sleep scoring

#### Electrophysiological in vivo recordings

Data was acquired using the 128 Channel Neurophysiology Recording System (Tucker-Davis Technologies Inc., Alachua, FL, USA) and the electrophysiological recording software Synapse (Tucker-Davis Technologies Inc., Alachua, FL, USA), and saved on a local computer. EEG and EMG signals were continuously recorded, filtered between 0.1 - 100 Hz, and stored at a sampling rate of 305 Hz. Extracellular neuronal activity was continuously recorded at a sampling rate of 25 kHz and filtered between 300 Hz - 5 kHz. Whenever the recorded voltage in an individual laminar channel crossed a manually set threshold indicating putative neuronal firing (at least 2 standard deviations above noise level), 46 samples around the event (0.48 ms before, 1.36 ms after) were stored. Concomitantly with the spike acquisition, LFPs were continuously recorded from the same electrodes and processed with the abovementioned settings for EEG signals.

#### Offline signal processing

EEG, EMG, and LFP signals were resampled at a sampling rate of 256 Hz using custom-made code in Matlab (The MathWorks Inc, Natick, Massachusetts, USA, version v2017a) and converted into the European Data Format (EDF) as previously^42^. Spike wave forms were further processed using a custom-made Matlab script and events with artefactual wave forms were excluded from further analysis of neuronal activity.

#### Scoring of vigilance states

The software Sleep Sign for Animals (SleepSign Kissei Comtec Co., Ltd., Nagano, Japan) was used for sleep scoring. EEG, EMG, and LFP recordings were partitioned into epochs of 4 s. Vigilance states were assigned manually to each recording epoch based on visual inspection of the frontal and occipital EEG derivations in conjunction with the EMG. For two animals with defective EEG reference, a fronto-occipital EEG derivation was used for sleep scoring. Epochs with recording artefacts due to gross movements, chewing or external electrostatic noise were assigned to the respective vigilance state but not included in the electrophysiological analysis. Overall 18.8±3.5 % of wake, 0.7±0.4 % of NREM, and 0.9±0.4 % of REM epochs contained artefactual EEG signals, with no significant difference between genotypes. EEG and LFP power spectra were computed using a fast Fourier transform routine (Hanning window) with a 0.25Hz resolution and exported in the frequency range between 0 and 30 Hz for spectral analysis.

### Statistical analysis

Data were analysed using MATLAB (version R2017b; The MathWorks Inc, Natick, Massachusetts, USA) and IBM SPSS Statistics for Windows, version 25.0 (IBM Corp., Armonk, N.Y., USA). Reported averages are mean±s.e.m. For analysis of repeated measures with more than 20 time or frequency bins (i.e. time courses of firing rates, latency to first spike for different spike numbers, and comparison of EEG power spectra), we performed unpaired t-tests for each individual time or frequency bin. Bins for which the resulting p-value was below 0.05 are highlighted with asterisks to indicate areas of interest bearing in mind that no correction for multiple comparisons was applied. EEG power spectra of individual animals were log-transformed before hypothesis testing. For comparison of time course data with less than 20 bins (i.e. time course of SWA after sleep deprivation or time course of time spent in NREM sleep), mixed ANOVAs with the within-subject factor ‘time interval’ and the between-subject factor ‘genotype’ or repeated-measure ANOVAs with the within-subject factors ‘time-interval’ and ‘experimental condition’ were applied and post-hoc comparisons were made using two-tailed unpaired or paired t-tests on individual time bins. Greenhouse-Geisser correction was used when the assumption of sphericity was violated (Mauchly’s test of sphericity, p<0.05). For the comparison of vigilance state duration between genotypes, we performed two-tailed t-tests or alternatively Mann-Whitney U tests if the assumption of normality was violated (Shapiro-Wilk test of normality, p<0.05). The NREM bout duration was compared between genotypes and time points using a mixed ANOVA with the within-subject factor ‘experimental condition’ and the between-subject factor ‘genotype’. Post-hoc comparisons were performed using two-tailed paired t-tests to compare time points for each genotype and using two-tailed unpaired t-tests to compare genotypes at each time point. In all figures, significance levels are indicated with asterisks: ‘*’ for 0.05 > p > 0.01; ‘**’ for 0.01 > p > 0.001; ‘***’ for 0.001 > p. Panels representing grouped data of durations spent in specific vigilance states show group mean (red line), 95% confidence interval (pink box), and one standard deviation (blue box) with individual data points overlaid and were generated using the MATLAB function notBoxPlot (Rob Campbell (2019). notBoxPlot (https://www.github.com/raacampbell/notBoxPlot), GitHub. Retrieved June 15, 2019).

### Criteria for analysis of vigilance state episodes and OFF periods

For analyses of mean and maximum duration of sustained wake episodes (Figure 2e), we included wake episodes, which were at least 1 min long allowing brief intrusions of sleep of 1 min or less. For the analysis of mean duration of NREM episodes, we included NREM episodes, which were at least 1 min long allowing brief intrusions of REM sleep or brief awakenings of 1 min or less (Figure 3e). To investigate the change in NREM SWA across prolonged wake episodes, we included consolidated periods of waking lasting at least 15 min, whereby short episodes of sleep <1 min were not considered as interruptions. We then performed analyses of NREM sleep SWA in the 15-min time window immediately preceding and following prolonged (>15 min) wake episodes, if both time windows included at least 10 min of artefact-free NREM sleep and no more than 3 min of wakefulness (Figure 2f). Population OFF periods were defined as periods of total neuronal silence across all electrodes, which lasted at least 50 ms and no more than 4000 ms. Subsequently, the top 20% longest OFF periods were included for final analyses (Fig 1f,g). The latency to the first spike after the population OFF-ON transition was calculated separately for MUA recorded in layers 2/3 and layer 5. Only ON periods with at least 1 spike in each of the layers occurring within the first 200 ms were included in this analysis (Figure 1g).

### Histological assessment of laminar probe depth

The tips of laminar implants were stained before surgery with the orange-red fluorescent^®^ membrane stain DiI (1,1’-Dioctadecyl-3,3,3’,3’-Tetramethylindocarbocyanine Perchlorate; Thermo Fisher Scientific, Waltham, MA, USA) for later histological assessment of the electrode position. After completion of the experiments, microlesions of selected channels on the laminar probe were performed under terminal anaesthesia using the electroplating device NanoZ™ (White Matter LLC, Seattle, WA, USA) applying 10 mA direct current for 25 s to each respective channel. Immediately following microlesioning, mice were perfused with 0.1M phosphate buffered saline (0.9 %) followed by 4 % paraformaldehyde for tissue preservation. Vibrating microtome (Leica VT1000S) sectioning was used to acquire 50 μm coronal brain slices. Fluorescent staining was performed with 4′,6-diamidino-2-phenylindole (DAPI). After fluorescence microscopy, implantation sites were mapped using a mouse brain atlas^43^ and the depth of the laminar implant was assessed measuring the distance between cortical surface and the electrical current induced tissue microlesions. ImageJ was used to merge fluorescent images and add scale bars^44^.

### Ethical approval

All experiments were performed in accordance with the United Kingdom Animal Scientific Procedures Act 1986 under personal and project licences granted by the United Kingdom Home Office. Ethical approval was provided by the Ethical Review Panel at the University of Oxford. Animal holding and experimentation were located at the Biomedical Sciences Building (BSB) and the Behavioural Neuroscience Unit (BNU), University of Oxford.

## Code availability

Custom-made code for data analysis used in this study is available from the corresponding authors upon request.

## Data availability

The datasets acquired for this study are available from the corresponding authors upon request.

## Acknowledgements

We thank Thomas Jahans-Price and Xudong Wang for their help with establishing the microlesion protocol; Merima Sabanovic, Vincent van der Vinne and Laura McKillop for their advice on statistics and figure production; Kristina Parley for her support in the histological procedures; Stuart Peirson for his advice on the experimental design; all members of the Vyazovskiy lab for their kind help with surgery assistance, animal care, and sleep deprivation. This work was supported by the Wellcome Trust PhD studentship 203971/Z/16/Z, Wellcome Trust Strategic Award 098461/Z/12/Z, John Fell OUP Research Fund Grant 131/032, and Medical Research Council (UK) grant MR/N026039/1.

## Author contributions

ZM and VVV initiated and proposed the study with pilot experiments done by TY. LBK, TY, CJA, AHS, ZM, and VVV designed the experiments. LBK, TY, MCCG, and CBD conducted the experiments on the transgenic mouse model. LBK, CBD, and MCK conducted the experiments on wild type mice. LBK, MCK, and AHS performed the histology. AHS and ZM developed, validated, and provided the transgenic mice. LBK and VVV analysed the data. LBK and VVV wrote the manuscript. All of the authors discussed the results and commented on the manuscript.

## Supplementary figures

**Suppl. Figure 1:**
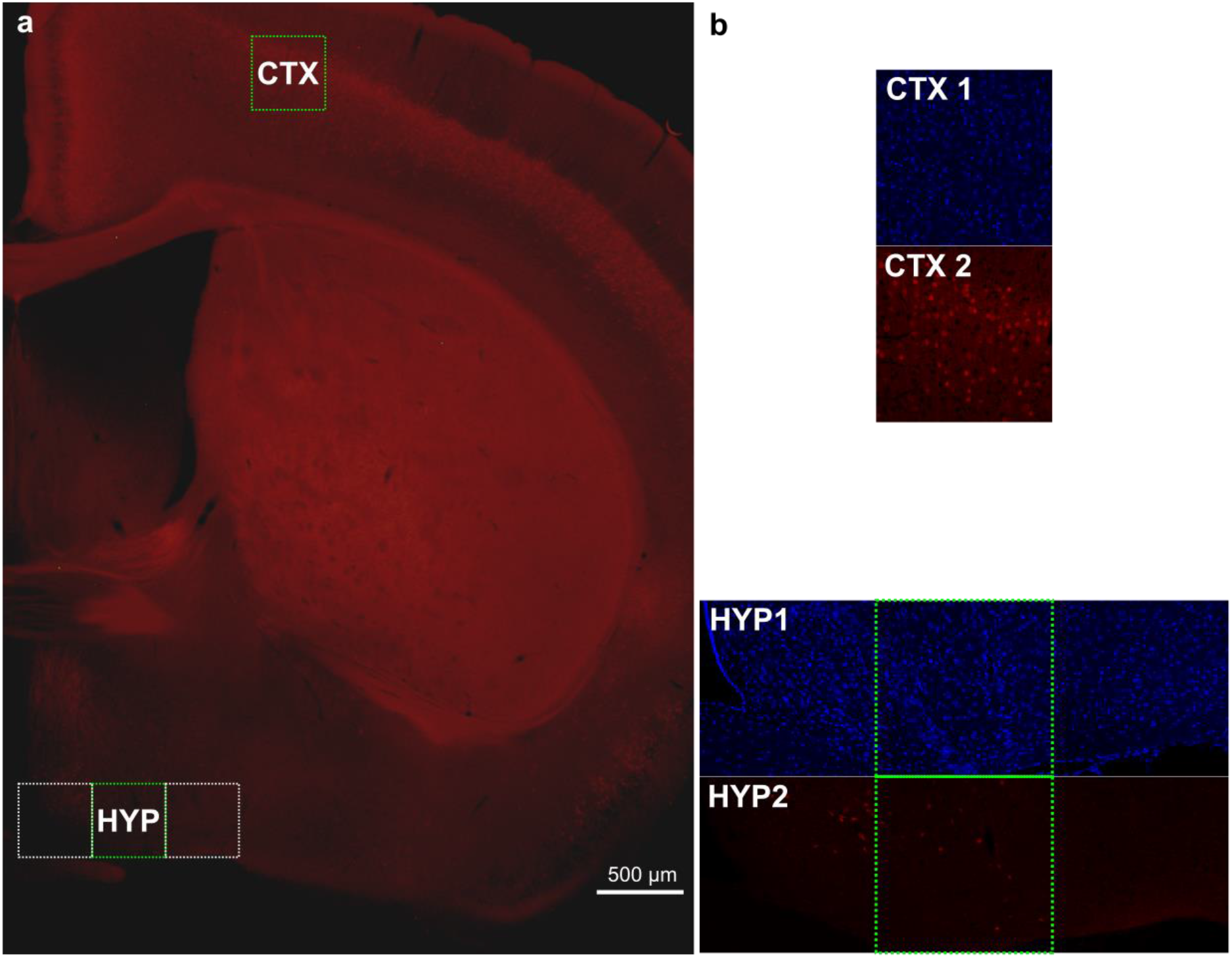
Comparison of Cre-expression in the Rbp4-Cre mouse line between cortical layer 5 and hypothalamus. a) Coronal section of a Rbp4-Cre;Ai14;Snap25fl/+ mouse brain (anterior-posterior position approximately Bregma) indicating areas, which were further examined for Cre-expression using confocal imaging of DAPI stained slices. For the hypothalamic area three neighbouring confocal tiles (425×425 μm each) were acquired starting from the lateral surface of the third ventricle. Cells were counted in the green highlighted rectangles of equal size. b) DAPI stained confocal images from cortex (CTX) and hypothalamus (HYP) in the blue fluorescence channel used for cell counting (CTX1 and HYP1) and in the red fluorescence channel (CTX2 and HYP2) used to identify Cre-expression in the respective cells indicated by expression of the red fluorescent STOP-floxed tdTomato reporter Ai14. Cell counts on corresponding coronal sections in three brains revealed that 20.53±0.98% (480/2342) of cortical cells were tdTomato positive, while only 1.15±0.40% (35/3006) of hypothalamic cells expressed the red fluorescent indicator.

**Suppl. Figure 2:**
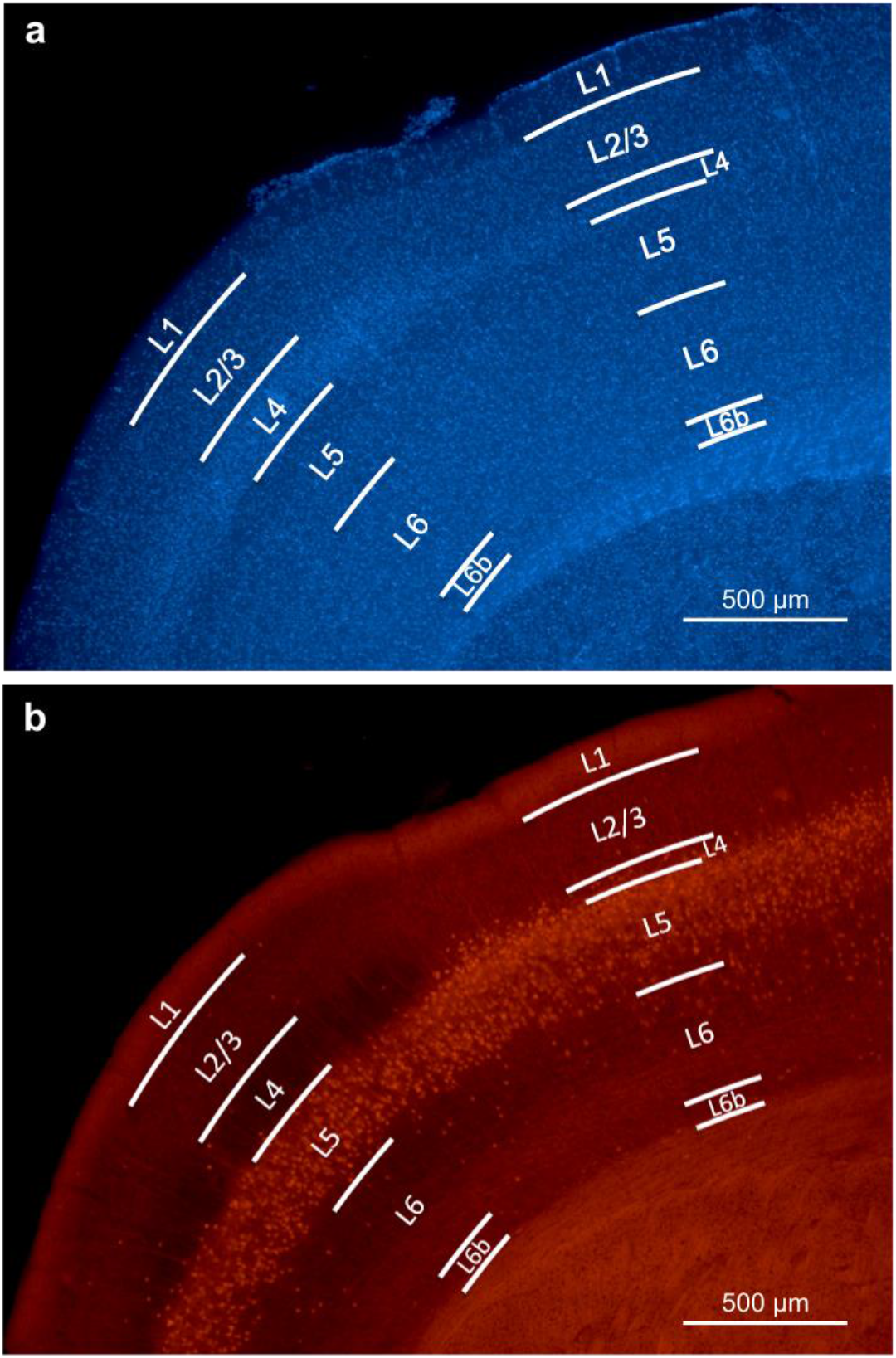
Identification of cortical layers in a representative coronal brain slice covering primary motor and sensory cortex. a) Cortical layers were determined using DAPI staining and identification of characteristic anatomical features of specific layers such as cell density and nuclear size. b) Image shows the largely restricted expression of the red fluorescent protein tdTomato in layer 5 in both primary motor and sensory cortex. The selective Cre-expression was driven by a Rbp4 promoter that cleaved the STOP-floxed site in the tdTomato reporter Ai14 mouse. Anterior-posterior position: approximately Bregma +0.75 mm.

**Suppl. Figure 3:**
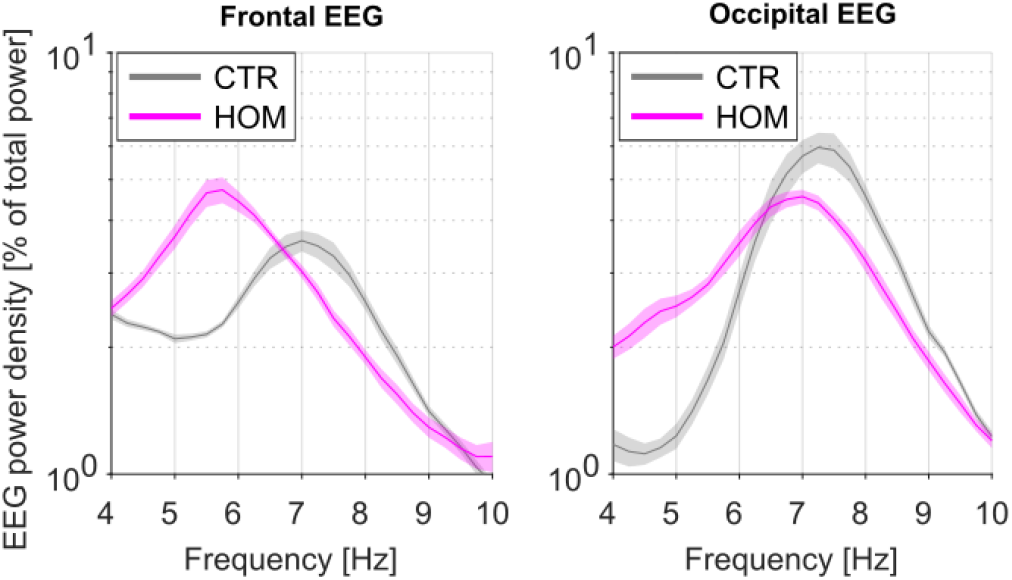
Relative spectral power in the theta frequency range during REM sleep in homozygous (HOM) Rbp4-Cre;Ai14;Snap25^fl/fl^ mice compared to Cre-negative littermates (CTR). Note that the peak of theta activity is shifted towards lower frequencies in HOMs (n=8) compared to CTRs (n=5) in both the frontal (HOM: 6.09±0.07 Hz; CTR: 6.95±0.09 Hz; p=0.002, Mann-Whitney U test) and occipital (HOM: 6.91±0.10 Hz; CTR: 7.35±0.06 Hz; p=0.015, Mann-Whitney U test) EEG derivations.

**Suppl. Figure 4:**
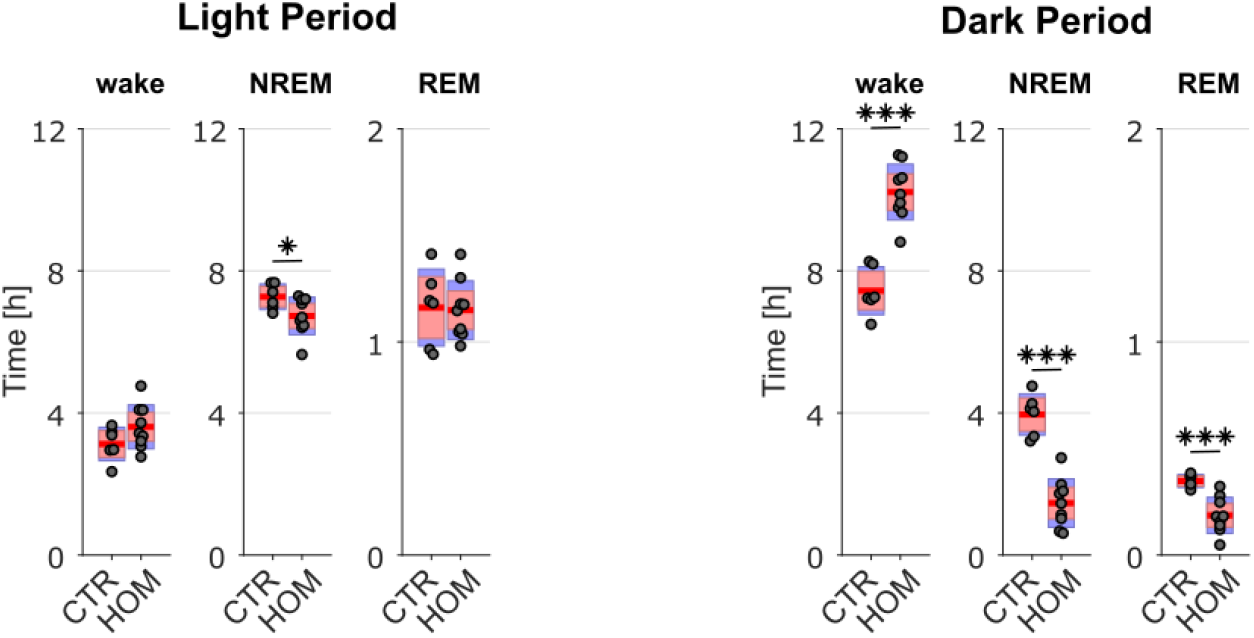
Amount of time spent in wake, NREM, and REM during the light period and dark period of undisturbed baseline recordings. During the light period, the distribution between vigilance states is similar between genotypes (wake: t(13)=−1.62, p=0.130; NREM: t(13)=2.17, p=0.049; REM: t(13)=0.149, p=0.884), while strong differences occur during the dark period (wake: t(13)=−7.07, p<0.001; NREM: t(13)=7.32, p<0.001; REM: t(13)=4.43, p=0.001). N=6 CTR and n=9 HOM. Data is presented as group mean (red line), 95% confidence interval (pink box), and one standard deviation (blue box) with individual data points overlaid.

**Suppl. Figure 5:**
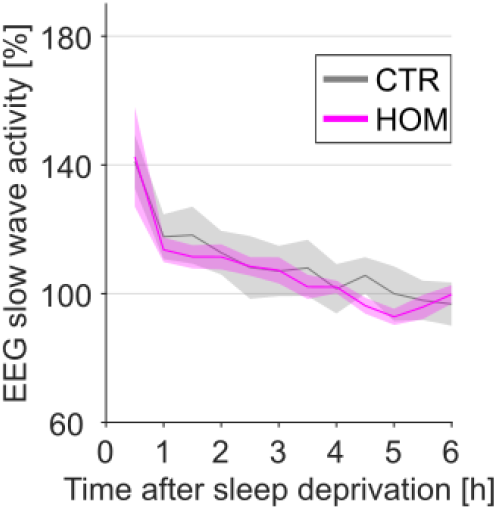
Time course of NREM slow wave activity (SWA) after sleep deprivation in the occipital EEG derivation. Time course of EEG slow wave activity during NREM sleep after sleep deprivation relative to baseline average. No significant differences between genotypes were observed (mixed ANOVA, within-subject factor ‘time’, between-subject factor ‘genotype’, main effect: F(1.907,20.975)=0.301, p=0.73. Post-hoc comparisons with two-tailed t-tests for independent samples reveals no significant differences for any of the 12 time intervals). N=5 CTR and n=8 HOM.

## References

1. Economo, C. V. Sleep as a problem of localization. J. Nerv. Ment. Dis. 71, 249–259 (1930).

2. Nauta, W. J. H. Hypothalamic regulation of sleep in rats; an experimental study. J. Neurophysiol. 9, 285–316 (1946).

3. Saper, C. B. & Fuller, P. M. Wake–sleep circuitry: an overview. Curr. Opin. Neurobiol. 44, 186–192 (2017).

4. Liu, D. & Dan, Y. A Motor Theory of Sleep-Wake Control: Arousal-Action Circuit. Annu. Rev. Neurosci. 42, 27–46 (2019).

5. Adamantidis, A. R., Gutierrez Herrera, C. & Gent, T. C. Oscillating circuitries in the sleeping brain. Nat. Rev. Neurosci. 20, 746–762 (2019).

6. Borbély, A. A. A two process model of sleep regulation. Hum. Neurobiol. 1, 195–204 (1982).

7. Vyazovskiy, V. V. et al. Cortical Firing and Sleep Homeostasis. Neuron 63, 865–878 (2009).

8. Vyazovskiy, V. V. & Harris, K. D. Sleep and the single neuron: the role of global slow oscillations in individual cell rest. Nat. Rev. Neurosci. 14, 443–451 (2013).

9. Tononi, G. & Cirelli, C. Sleep and the Price of Plasticity: From Synaptic and Cellular Homeostasis to Memory Consolidation and Integration. Neuron 81, 12–34 (2014).

10. Huber, R., Felice Ghilardi, M., Massimini, M. & Tononi, G. Local sleep and learning. Nature 430, 78–81 (2004).

11. Vyazovskiy, V. V. et al. Local sleep in awake rats. Nature 472, 443–7 (2011).

12. Krueger, J. M., Nguyen, J. T., Dykstra-Aiello, C. J. & Taishi, P. Local sleep. Sleep Med. Rev. 43, 14–21 (2019).

13. Sanchez-Vives, M. V & Mattia, M. Slow wave activity as the default mode of the cerebral cortex. Arch Ital Biol 152, 147–155 (2014).

14. González-Rueda, A., Pedrosa, V., Feord, R. C., Clopath, C. & Paulsen, O. Activity-Dependent Downscaling of Subthreshold Synaptic Inputs during Slow-Wave-Sleep-like Activity In Vivo. Neuron 97, 1244–1252.e5 (2018).

15. Lemieux, M., Chen, J.-Y., Lonjers, P., Bazhenov, M. & Timofeev, I. The Impact of Cortical Deafferentation on the Neocortical Slow Oscillation. J. Neurosci. 34, 5689 LP – 5703 (2014).

16. Saberi-Moghadam, S., Simi, A., Setareh, H., Mikhail, C. & Tafti, M. In vitro Cortical Network Firing is Homeostatically Regulated: A Model for Sleep Regulation. Sci. Rep. 8, 6297 (2018).

17. Neske, G. T. The Slow Oscillation in Cortical and Thalamic Networks: Mechanisms and Functions. Front. Neural Circuits 9, 88 (2016).

18. van Aerde, K. I., Qi, G. & Feldmeyer, D. Cell Type-Specific Effects of Adenosine on Cortical Neurons. Cereb. Cortex 25, 772–787 (2015).

19. Lőrincz, M. L. et al. A distinct class of slow (~0.2-2 Hz) intrinsically bursting layer 5 pyramidal neurons determines UP/DOWN state dynamics in the neocortex. J. Neurosci. 35, 5442–58 (2015).

20. Beltramo, R. et al. Layer-specific excitatory circuits differentially control recurrent network dynamics in the neocortex. Nat. Neurosci. 16, 227–234 (2013).

21. Chauvette, S., Volgushev, M. & Timofeev, I. Origin of Active States in Local Neocortical Networks during Slow Sleep Oscillation. Cereb. Cortex 20, 2660–2674 (2010).

22. Sanchez-Vives, M. V & McCormick, D. A. Cellular and network mechanisms of rhytmic recurrent activity in neocortex. Nat. Neurosci. 3, 1027–1034 (2000).

23. Sakata, S. & Harris, K. D. Laminar Structure of Spontaneous and Sensory-Evoked Population Activity in Auditory Cortex. Neuron 64, 404–418 (2009).

24. Funk, C. M., Honjoh, S., Rodriguez, A. V., Cirelli, C. & Tononi, G. Local slow waves in superficial layers of primary cortical areas during REM sleep. Curr. Biol. 26, 396–403 (2016).

25. Hoerder-Suabedissen, A. et al. Cell-Specific Loss of SNAP25 from Cortical Projection Neurons Allows Normal Development but Causes Subsequent Neurodegeneration. Cereb. Cortex 29, 2148–2159 (2019).

26. Gerfen, C. R., Paletzki, R. & Heintz, N. GENSAT BAC cre-recombinase driver lines to study the functional organization of cerebral cortical and basal ganglia circuits. Neuron 80, 1368–1383 (2013).

27. Washbourne, P. et al. Genetic ablation of the t-SNARE SNAP-25 distinguishes mechanisms of neuroexocytosis. Nat. Neurosci. 5, 19–26 (2002).

28. Buzsáki, G. Theta Oscillations in the Hippocampus. Neuron 33, 325–340 (2002).

29. Boyce, R., Glasgow, S. D., Williams, S. & Adamantidis, A. Causal evidence for the role of REM sleep theta rhythm in contextual memory consolidation. Science (80-.). 352, 812 LP – 816 (2016).

30. Huber, R., Deboer, T. & Tobler, I. Effects of sleep deprivation on sleep and sleep EEG in three mouse strains: empirical data and simulations. Brain Res. 857, 8–19 (2000).

31. Vyazovskiy, V. V, Ruijgrok, G., Deboer, T. & Tobler, I. Running Wheel Accessibility Affects the Regional Electroencephalogram during Sleep in Mice. Cereb. Cortex 16, 328–336 (2005).

32. Huber, R., Deboer, T. & Tobler, I. Topography of EEG Dynamics After Sleep Deprivation in Mice. J. Neurophysiol. 84, 1888–1893 (2000).

33. Franken, P., Dijk, D. J., Tobler, I. & Borbely, A. A. Sleep deprivation in rats: effects on EEG power spectra, vigilance states, and cortical temperature. Am. J. Physiol. Integr. Comp. Physiol. 261, R198–R208 (1991).

34. Frank, M. G. & Heller, H. C. The Function(s) of Sleep. Handbook of Experimental Pharmacology 253, 3–34 (2019).

35. Morairty, S. R. et al. A role for cortical nNOS/NK1 neurons in coupling homeostatic sleep drive to EEG slow wave activity. Proc Natl Acad Sci U S A 110, 20272–20277 (2013).

36. Eichenbaum, H. On the Integration of Space, Time, and Memory. Neuron 95, 1007–1018 (2017).

37. Greene, R. W., Bjorness, T. E. & Suzuki, A. The adenosine-mediated, neuronal-glial, homeostatic sleep response. Curr. Opin. Neurobiol. 44, 236–242 (2017).

38. Brüning, F. et al. Sleep-wake cycles drive daily dynamics of synaptic phosphorylation. Science (80-.). 366, eaav3617 (2019).

39. Honda, T. et al. A single phosphorylation site of SIK3 regulates daily sleep amounts and sleep need in mice. Proc. Natl. Acad. Sci. 115, 10458 LP – 10463 (2018).

40. Williams, J. A. & Naidoo, N. Sleep and cellular stress. Curr. Opin. Physiol. 15, 104–110 (2020).

41. Kempf, A., Song, S. M., Talbot, C. B. & Miesenböck, G. A potassium channel β-subunit couples mitochondrial electron transport to sleep. Nature 568, 230–234 (2019).

42. McKillop, L. E. et al. Effects of Aging on Cortical Neural Dynamics and Local Sleep Homeostasis in Mice. J. Neurosci. 38, 3911 LP – 3928 (2018).

43. Paxinos, G. & Franklin, K. B. J. Paxinos and Franklin’s the mouse brain in stereotaxic coordinates. (Elsevier Academic Press, 2019).

44. Schindelin, J. et al. Fiji: an open source platform for biological image analysis. Nat. Methods 9, 676–682 (2012).

